# Accurate and Efficient KIR Gene and Haplotype Inference from Genome Sequencing Reads with Novel K-mer Signatures

**DOI:** 10.1101/541938

**Authors:** David Roe, Rui Kuang

## Abstract

The killer cell immunoglobulin-like receptor (KIR) proteins evolve to fight viruses and mediate the body’s reaction to pregnancy. These roles provide selection pressure for variation at both the structural/haplotype and base/allele levels. At the same time, the genes have evolved relatively recently by tandem duplication and therefore exhibit very high sequence similarity over thousands of bases. These variation-homology patterns make it impossible to interpret KIR haplotypes from abundant short-read genome sequencing data at population scale using existing methods. Here, we developed an efficient computational approach for *in silico* KIR probe interpretation (KPI) to accurately interpret individual’s KIR genes and haplotype-pairs from KIR sequencing reads. We designed synthetic 25-base sequence probes by analyzing previously reported haplotype sequences, and we developed a bioinformatics pipeline to interpret the probes in the context of 16 KIR genes and 16 haplotype structures. We demonstrated its accuracy on a synthetic data set as well as a real whole genome sequences from 748 individuals from The Genome of the Netherlands (GoNL). The GoNL predictions were compared with predictions from SNP-based predictions. Our results show 100% accuracy rate for the synthetic tests and a 99.6% family-consistency rate in the GoNL tests. Agreement with the SNP-based calls on KIR genes ranges from 72-100% with a mean of 92%; most differences occur in genes *KIR2DS2, KIR2DL2, KIR2DS3*, and *KIR2DL5* where KPI predicts presence and the SNP-based interpretation predicts absence. Overall, the evidence suggests that KPI’s accuracy is 97% or greater for both KIR gene and haplotype-pair predictions, although the presence/absence genotyping leads to ambiguous haplotype-pair predictions with 16 reference KIR haplotype structures. KPI is free, open, and easily executable as a Nextflow workflow supported by a Docker environment at https://github.com/droeatumn/kpi.

## Introduction

Human chromosome 19q13.4 contains a ∼150-250 kilobase region encoding 16 genes of the natural killer cell immunoglobulin-like receptor (KIR) family. These genes are ∼4-16 kilobases long and evolved via tandem duplication during primate evolution(1)(2). The KIR receptors recognize human leukocyte antigen (HLA) class I molecules and contribute to natural killer (NK) cell functions via activating or inhibiting signals. These receptor-ligand pairs coevolved under selection pressures from reproduction and pathogenic defense(3), and it is believed that KIR genes have undergone a balancing selection via duplications and deletions into two broad categories of haplotypes, in which one category tends to vary more at the allelic level and the other tends to vary more at the structural (gene content and order) level(4)(5)(6). A few dozen KIR full haplotype sequences and approximately 2500 full- or inter-gene sequences have been publicly deposited(7)(5)(8). Haplotype structures are divided into two classes. Class ‘A’ contains one haplotype and its deleted forms. Class ‘B’ haplotypes are more structurally diverse and contain a variety of insertions and deletions. Generally, the A haplotype occurs with 50-60% frequency, haplotypes that are half-A and half-B occur with 30-40%, and the rest of the haplotypes are variants of the B haplotypes. Except for some rarer deleted forms, KIR haplotypes are structurally variable around 4 ‘framework’ genes (*KIR3DL3, KIR3DP1, KIR2DL4, KIR3DL2*), with *KIR3DL3* through *KIR3DP1* defining the proximal (or ‘centromeric’) region and *KIR2DL4* through *KIR3DL2* defining the distal (or ‘telomeric’) region, with the two gene-rich regions separated by the relatively large and recombinant *KIR3DP1-KIR2DL4* intergene region.

It is difficult to interpret the KIR region with high-throughput sequencing reads for an individual human genome when the structural arrangements are unknown; indeed, it is difficult even when the structural haplotypes *are* known, since the read length is too short to map unambiguously to the repetitive and homologous KIR genes. As a consequence, the reads from KIR region are largely ignored, mis-interpreted, or under-interpreted in current whole genome sequencing (WGS) studies. SNP (single nucleotide polymorphism)-based KIR interpretation is more commonly applied. For example, KIR*IMP is a web-application to predict genes and haplotypes from microarray SNP genotypes(9). As an algorithm whose raw data is microarray calls, KIR*IMP can interpret KIR from genome wide SNP arrays, but it is not applicable to interpret KIR from raw sequences.

Since a general solution for KIR structural interpretation from raw genomic DNA is not currently available, this study implements such an algorithm for the prediction of KIR genes and full structural haplotypes from any type of raw full-region-or-greater genomic sequence at population scale. In particular, we systematically evaluated small markers for KIR genes and then applied those markers to a synthetic KIR probe interpretation (KPI) algorithm for the presence/absence of 16 KIR genes and 16 haplotype structures. Our approach leverages recent bioinformatics innovations for short sequence (‘probe’) genotyping, along recently published KIR reference haplotypes. The KPI algorithm first efficiently counts the occurrence of each kmer probe in the raw sequences, and then uses multiple probes per gene to call its presence/absence. Those 16 genotypes are then used to generate haplotype-pair predictions. In the experiments, we report 100% accuracy on a test set of synthetic haplotypes for comparisons with known truth. We also report that gene and haplotype-pair predictions for the WGS GoNL cohort are family consistent and compare favorably with reference frequencies in comparison to SNP-based predictions using KIR*IMP.

## Materials and methods

### Overview

The workflow of KPI consists of three steps,

1. Discover the 25mer gene markers based on a multiple sequence alignment analysis of 68 full-length haplotype sequences.
2. Count the 25mer markers in the reads of genomic DNA per individual to generate the individual’s 25mer genotype.
3. Predict presence/absence per gene from the marker genotypes for each individual.
4. Predict haplotype pairs from the gene presence/absence calls for each individual.

In the following, we first explain each step and then describe the synthetic data and GoNL data used for the evaluation.

### Step 1: Discovering 25mer gene markers

To discover gene marker 25mers, first a multiple sequence alignment (MSA) was created with 68 publicly deposited full-length haplotypes sequences (manuscript under preparation). Briefly, each haplotype was annotated at an average resolution of ∼4kbp using a set of 15 120-base markers. This high-level annotation was aligned into a MSA representing a structural alignment of all haplotypes. Then, each subregion was aligned to base pair resolution. This resulted in a full resolution, full haplotype MSA that accurately classifies each allele into a haplotype-defined locus, and it aligns the alleles precisely at each locus. The haplotype and gene annotations of the MSA provided a list of full-length alleles for 16 genes: *KIR2DL1-5, KIR2DS1-5, KIR2DP1, KIR3DL1-3, KIR3DP1*, and *KIR3DS1*. Markers for each gene locus were chosen by selecting all sequences of length 25 (25mers) present in every allele of the gene but not elsewhere in the KIR haplotypes nor the rest of the genome reference GRCh38 (details in Supplementary Figure 1).

### Step 2: Count 25mer markers

KMC 3, with workflows implemented in Nextflow(13) and Apache Groovy(14) and a software environment implemented as a Docker container, is used to create 25mer databases from sequence or short-read data and match the markers across the datasets. Using KMC 3, we generate the list of all 25mers from the short reads of each individual and then match the 25mers in the marker databases to report the hit counts of each 25mer marker in the individual. Details are in Supplemental Figure 1.

### Step 3: Individual genotyping from 25mer markers

KPI calls presence/absence per gene by aggregating the presence/absence genotypes of many small (25mer) markers, each specific to one gene. 25mers with hit counts less than three are considered sequencing errors and set to zero. If the mean hit count of all the markers per gene is zero, then the gene is predicted absent; otherwise, it is called present. Additional details can be found in Supplemental Figure 1.

### Step 4: Individual haplotyping from genotypes

Haplotype-pair predictions were made by fitting the genotype to all possible pairs of the 16 structural reference haplotypes defined in Table 1. The numbers and frequencies of the haplotypes are from Jiang et al. 2012(4); some of their haplotypes are combined in Table 1 because they consider certain alleles as separate haplotypes, such as full or deleted alleles of *KIR2DS4*. These 16 haplotypes represent 97% of all haplotypes in the Jiang et al. report.

**Table 1.** Reference haplotype definitions. Haplotype numbers (Jiang et al. 2012) and informal names are shown with a reference frequency and their definition via gene counts. Following Jiang et al. convention, some haplotypes (e.g., 7, 9) are distinguished by *KIR2DS3*/*KIR2DS5* alleles instead of structural differences. In this study, some haplotypes (e.g., 1, 2) are combined, as *KIR2DS4* full/deleted alleles are not considered in KPI’s genotyping.

| Jiang et al. 2012 # | informal nomenclature | Jiang et al. 2012 freq. | 3DL3 | 2DS2 | 2DL2 | 2DL3 | 2DP1 | 2DL1 | 3DP1 | 2DL4 | 3DL1 | 3DS1 | 2DL5 | 2DS3 | 2DS5 | 2DS4 | 2DS1 | 3DL2 |
| --- | --- | --- | --- | --- | --- | --- | --- | --- | --- | --- | --- | --- | --- | --- | --- | --- | --- | --- |
| 1/2 | cA01~tA01 | 55.2% | 1 | 0 | 0 | 1 | 1 | 1 | 1 | 1 | 1 | 0 | 0 | 0 | 0 | 1 | 0 | 1 |
| 3 | cA01~tB01_2DS5 | 10.9% | 1 | 0 | 0 | 1 | 1 | 1 | 1 | 1 | 0 | 1 | 1 | 0 | 1 | 0 | 1 | 1 |
| 11 | cA01~tB01_2DS3 | 1.4% | 1 | 0 | 0 | 1 | 1 | 1 | 1 | 1 | 0 | 1 | 1 | 1 | 0 | 0 | 1 | 1 |
| 4/5 | cB02~tA01 | 12.8% | 1 | 1 | 1 | 0 | 0 | 0 | 1 | 1 | 1 | 0 | 0 | 0 | 0 | 1 | 0 | 1 |
| 6/10 | cB01~tA01_2DS3 | 6.9% | 1 | 1 | 1 | 0 | 1 | 1 | 1 | 1 | 1 | 0 | 1 | 1 | 0 | 1 | 0 | 1 |
| 25 | cB01~tA01_2DS5 | 0.1% | 1 | 1 | 1 | 0 | 1 | 1 | 1 | 1 | 1 | 0 | 1 | 0 | 1 | 1 | 0 | 1 |
| 7 | cB01~tB01_2DS3_2DS5 | 2.6% | 1 | 1 | 1 | 0 | 1 | 1 | 1 | 1 | 0 | 1 | 1 | 1 | 1 | 0 | 1 | 1 |
| 9 | cB01~tB01_2DS3_2DS3 | 2.1% | 1 | 1 | 1 | 0 | 1 | 1 | 1 | 1 | 0 | 1 | 1 | 2 | 0 | 0 | 1 | 1 |
| 8 | cB02~tB01_2DS5 | 2.1% | 1 | 1 | 1 | 0 | 0 | 0 | 1 | 1 | 0 | 1 | 1 | 0 | 1 | 0 | 1 | 1 |
| 17 | cB02~tB01_2DS3 | 0.3% | 1 | 1 | 1 | 0 | 0 | 0 | 1 | 1 | 0 | 1 | 1 | 1 | 0 | 0 | 1 | 1 |
| 12 | cB04~tB03_2DS5 | 0.8% | 1 | 1 | 1 | 0 | 0 | 0 | 0 | 0 | 0 | 1 | 1 | 0 | 1 | 0 | 1 | 1 |
| 18 | cB04~tB03_2DS3 | 0.3% | 1 | 1 | 1 | 0 | 0 | 0 | 0 | 0 | 0 | 1 | 1 | 1 | 0 | 0 | 1 | 1 |
| 13 | cB01~tB05 | 0.7% | 1 | 1 | 1 | 0 | 1 | 1 | 2 | 2 | 1 | 1 | 1 | 1 | 0 | 1 | 0 | 1 |
| 15 | cB05~tB01 | 0.4% | 1 | 1 | 0 | 0 | 1 | 1 | 1 | 1 | 0 | 1 | 1 | 0 | 1 | 0 | 1 | 1 |
| 16 | cA01~tB05 | 0.3% | 1 | 0 | 0 | 1 | 1 | 1 | 2 | 2 | 1 | 1 | 0 | 0 | 0 | 1 | 0 | 1 |
| 21 | cB05~tA01 | 0.2% | 1 | 1 | 0 | 0 | 1 | 1 | 1 | 1 | 1 | 0 | 0 | 0 | 0 | 1 | 0 | 1 |
| sum |  | 97.0% |  |  |  |  |  |  |  |  |  |  |  |  |  |  |  |  |

For the GoNL predictions, haplotype ambiguity was reduced by family trio patterns and then further by the EM (Expectation-Maximization)-based methods as described and used in Vierra-Green 2012(10). Haplotype frequencies were calculated from the EM-reduced individual haplotype-pair predictions. These haplotype frequency calculations are not possible on the KPI’s haplotype-pair predictions because they can be ambiguous.

### Synthetic capture on diploid data

KPI was evaluated on a synthetic test set. There are six reference haplotype structures with publicly deposited full-length sequences (Table 1, top six rows). For each of these six structures, one sequence was randomly chosen to represent that structure, and it was paired with a random haplotype sequence from the set of all sequences. dwgsim(11) was used to generate 10,000 2×150 pair reads per haplotype (∼20x coverage) with 1% error rate. This provided a simulated six-person validation set of six diploid whole-region short-read sequences, representing all fully sequenced haplotype structures and paired to provide a variety of genotypes.

### GoNL family WGS and Immunochip SNP Data

KPI was also run on a large real-world example. WGS was obtained from The Genome of the Netherlands(GoNL)(12), a genome sequencing project whose goal is to map the genetic variation within the population of the Netherlands in 250 family trios (750 individuals). The project provided non-paired sequencing of the whole genomes of the population, which was done on the Illumina HiSeq 2000 platform. Coverage of the KIR region were similar to the previously reported(12) whole-genome average of ∼10-15x. Two individuals from two different families were removed from the GoNL project for data quality reasons, giving a total of 748 individuals.

KPI’s GoNL predictions were compared with results from microarray-based interpretation algorithm KIR*IMP. Illumina Immunochip microarray SNP data was obtained from GoNL(12). The data was prepared and executed following instructions using KIR*IMP v1.2.0 on 2019-10-05.

To the best of our knowledge, there only exists one method to predict KIR gene content from WGS sequences(13). However, we were unable to obtain results with it for both evaluation data sets. According to the authors, the current version is deprecated and to be replaced soon(14).

## Results

The predictions were evaluated in the small synthetic test set, where truth is known and a large real-world test set, where truth is unknown except for family relationships. Predictions were evaluated by comparing gene and haplotype-pair predictions to: known truth in the synthetic cohort, and family consistency (real-world cohort only), reference frequencies from Jiang et al.’s family copy number study(4), and the Allele Frequencies Database(15) in the real-world cohort. The real-world cohort was also compared with predictions from microarray-based algorithm KIR*IMP, although KIR*IMP was not considered ground truth as it reports accuracies as low as 81% for some genes(9). Haplotype-pair predictions were considered to be family consistent if each parents’ two haplotype predictions contained at least one of the child’s two predictions and one of the child’s haplotypes occurred in one parent and the other haplotype occurred in the other parent.

### Synthetic evaluation

Table 2 shows the results of the synthetic tests. The gene present/absent calls were 100% accurate for all genes. Although the haplotype predictions are ambiguous in half of the individuals, all are consistent with the ground truth.

**Table 2.** Results of synthetic tests. The first four columns detail the sequences from which the tests (n=6 haplotype-pairs) were generated. The fifth column is KPI’s haplotype predictions, some of which are summarized for display.

| haplotype 1 structure | haplotype 2 structure | haplotype 1 GenBank accession | haplotype 2 GenBank accession | KPI haplotype prediction | haplotypes consistent w/truth? | gene prediction accuracy |
| --- | --- | --- | --- | --- | --- | --- |
| cA01~tA01 | cA01~tA01 | GU182344 | GU182340 | cA01~tA01+cA01~tA01 | Y | 100% |
| cA01~tB01 | cA01~tA01 | KU645197 | GU182360 | cA01~tA01+cA01~tB01<br>or<br>cA01~tB01+cA01~tB05 | Y | 100% |
| cB01~tA01 | cA01~tA01 | GU182351 | NC000019.10 | cA01~tA01+cB01~tA01 | Y | 100% |
| cB01~tB01 | cB02~tA01 | GU182339 | GU182353 | 10 possibilities, including<br>cB01~tB01+cB02~tA01 | Y | 100% |
| cB02~tA01 | cA01~tA01 | GU182341 | KP420442 | cA01~tA01+cB02~tA01 | Y | 100% |
| cB02~tB01 | cA01~tA01 | GU182359 | KP420439 | 9 possibilities, including<br>cA01~tA01+cB02~tB01 | Y | 100% |

### GoNL evaluation

Table 3 shows a summary of the gene prediction results from KPI and KIR*IMP on the GoNL data set. A reference frequency range is included from Allele Frequencies Net, selecting European cohorts >= 500 individuals. Overall agreement between KIR*IMP and KPI for the 16 genes (Table 3, column 6) ranges from 72% to 100%, with a mean of 92%. KIR*IMP differs from the reference haplotype (Table 3, column 7) frequency range by >10% in 4 genes (*KIR2DS2, KIR2DL2, KIR2DL5, and KIR2DS3*) compared with 0 genes for KPI (Table 3, column 8). Both KIR*IMP and KPI differ from the KIR2DS1 reference by 9-10%, although the two algorithms agree in 98% of individuals for that gene.

**Table 3.** Summary of KIR*IMP and KPI gene predictions. Frequencies relative to GoNL cohort of 748 individuals. The abbreviated gene name is in column 1. Column 2 lists the reference frequencies from The Allele Frequency Net Database. The predicted frequencies from KIR*IMP and KPI are in columns 3 and 4, respectively. The delta between KIR*IMP and KPI is shown in the column 5. Column 6 shows the agreement between KIR*IMP and KPI. Column 7 shows the delta between KIR*IMP and the reference. Column 8 shows the delta between KPI and the reference. Frequencies with differences >10% are in bold.

| gene | reference freq. | KIR*IMP freq. | KPI freq. | KIR*IMP - KPI | KIR*IMP & KPI agreement | KIR*IMP - reference | KPI - reference |
| --- | --- | --- | --- | --- | --- | --- | --- |
| 2DS2 | 53-54% | 39% | 52% | -13% | **72%** | **-14%** | **-1%** |
| 2DL2 | 53-54% | 39% | 51% | -12% | **72%** | **-14%** | **-2%** |
| 2DL3 | 90% | 96% | 92% | 4% | 92% | 6% | 2% |
| 2DP1 | 96% | 99% | 97% | 2% | 97% | 3% | 1% |
| 2DL1 | 96% | 99% | 97% | 2% | 97% | 3% | 1% |
| 3DP1 | 100% | 100% | 100% | 0% | 100% | 0% | 0% |
| 2DL4 | 100% | 100% | 100% | 0% | 100% | 0% | 0% |
| 3DL1 | 93-94% | 96% | 96% | 0% | 100% | 2% | 2% |
| 3DS1 | 38-44% | 33% | 35% | -2% | 97% | -5% | -3% |
| 2DL5 | 53-56% | 38% | 47% | -9% | **87%** | **-15%** | **-6%** |
| 2DS3 | 30-31% | 10% | 29% | -18% | **81%** | **-20%** | **-1%** |
| 2DS5 | 30-36% | 25% | 27% | -2% | 96% | -5% | -3% |
| 2DS4 | 92-94% | 96% | 96% | 0% | 100% | 2% | 2% |
| 2DS1 | 43-44% | 33% | 34% | -1% | 98% | -10% | -9% |
| average |  |  |  |  | 92% |  |  |

Table 4 breaks down the differences between KIR*IMP and KPI in a confusion matrix. In the cases where KIR*IMP calls present (‘P’) and KPI calls absent (‘A’) (Table 4, column 2), the largest discrepancies are found in the centromeric genes *KIR2DS2* (8%), *KIR2DL2* (8%), and *KIR2DL3* (6%). In the reverse cases, when KIR*IMP calls absent and KPI calls present (Table 4, column 3), the largest discrepancies are greater and occur with the centromeric *KIR2DS2* (20%), *KIR2DL2* (20%), and the paralogous (centromeric or telomeric) *KIR2DL5* (11%), and *KIR2DS3* (19%).

**Table 4.** Confusion matrix of KIR*IMP and KPI gene predictions. Frequencies relative to GoNL cohort size of 748 individuals. The abbreviated gene names are in column 1. Column 2 lists the cases when KIR*IMP calls present (‘P’) and KPI calls absent (‘A’). Column 3 lists the cases when KIR*IMP calls absent (‘A’) and KPI calls present (‘P’). Column 4 is when they both call present. Column 5 is when they both call absent. Discrepancies >10% are in bold.

| gene | KIR*IMP: P<br>KPI: A | KIR*IMP: A<br>KPI: P | KIR*IMP: P<br>KPI: P | KIR*IMP: A<br>KPI: A |
| --- | --- | --- | --- | --- |
| 2DS2 | 8% | **20%** | 32% | 40% |
| 2DL2 | 8% | **20%** | 31% | 41% |
| 2DL3 | 6% | 2% | 90% | 2% |
| 2DP1 | 2% | 0% | 97% | 0% |
| 2DL1 | 2% | 1% | 97% | 0% |
| 3DP1 | 0% | 0% | 100% | 0% |
| 2DL4 | 0% | 0% | 100% | 0% |
| 3DL1 | 0% | 0% | 96% | 4% |
| 3DS1 | 1% | 3% | 32% | 64% |
| 2DL5 | 2% | **11%** | 36% | 51% |
| 2DS3 | 1% | **19%** | 10% | 71% |
| 2DS5 | 1% | 3% | 24% | 72% |
| 2DS4 | 0% | 0% | 96% | 4% |
| 2DS1 | 1% | 1% | 33% | 66% |

Per-individual haplotype-pair predictions for the GoNL cohort are included in Supplemental Table 1. Three lists of haplotype-pairs are provided: one for the initial fitting of all possible haplotype-pairs that could explain the genotype (i.e., KPI’s output); another that reduces those possibilities by family relationships; and one for the EM-reduced final haplotype-pair predictions.

KIR*IMP makes one most-likely prediction for all individuals. KPI’s predictions are sometimes ambiguous, with most predictions (mode) having one haplotype-pair but a mean of 2.3, standard deviation of 2.5, and a maximum of 14 haplotype-pair predictions per individual in the context of the 16 reference haplotypes. The KIR*IMP predictions are family consistent 100% of the time compared with 99.6% for KPI. However, the haplotype pair predictions between the two algorithms are concordant only 58% of the predictions.

Table 5 compares the haplotype predictions between KIR*IMP and the EM-reduced haplotype pair predictions from the KPI output. KIR*IMP fit 100% of its predicted genotypes into 15 of its reference haplotypes. KPI fit 97% of its predicted genotypes into its 16 reference haplotypes. KIR*IMP made predictions for two haplotypes (cA01∼tB04 and cB04∼tB03, numbered 14, 18, and 12), totaling 0.47%, that are not in KPI’s set of reference haplotypes. KPI’s haplotype-pair predictions are too ambiguous to summarize in haplotype frequencies.

**Table 5.** Comparison of KIR*IMP highest probability and EM-reduced haplotype prediction frequencies. The table shows the comparison of the predictions between both methods as well as with reference European frequencies from Jiang et al. 2012 (column 2), which is the source of the haplotype numbers (column 1). KIR*IMP’s haplotype frequencies for the 1496 GoNL haplotypes are in column 3; some haplotypes are combined, as the haplotype numbers distinguish KIR2DS4 alleles. Column 4 contains frequencies for EM-reduced KPI haplotype predictions Column 5 compares KIR*IMP frequencies with the reference, as column 6 does for EM predictions. Finally, column 7 compares the frequencies of KIR*IMP and the EM-reduced predictions. Haplotypes with a predicted frequency of 0 in both KIR*IMP and KPI are not shown. * Haplotypes 14, 18, and 12 are in KIR*IMP’s set of reference haplotypes, but not KPI’s.

| hap<br># | reference<br>frequency | KIR*IMP | KPI<br>w/EM | KIR*IMP<br>-<br>reference | KPI w/EM<br>-<br>reference | KIR*IMP<br>-<br>KPI<br>w/EM |
| --- | --- | --- | --- | --- | --- | --- |
| 1 | 55.20% | 71.86% | 59.70% | 16.70% | 4.50% | 12.17% |
| 2 |  |  |  |  |  |  |
| 3 | 10.90% | 12.57% | 9.60% | 1.70% | -1.29% | 2.95% |
| 11 | 1.40% | 1.47% | 0.60% | 0.00% | -0.82% | 0.85% |
| 4 | 12.80% | 7.49% | 15.30% | -5.30% | 2.55% | -7.83% |
| 5 |  |  |  |  |  |  |
| 9 | 2.10% | 3.41% | 2.90% | 1.30% | 0.78% | 0.52% |
| 7 | 2.60% | 0.33% | 3.60% | -2.20% | 1.00% | -3.24% |
| 6 | 6.90% | 1.80% | 5.90% | -5.10% | -1.08% | -4.05% |
| 10 |  |  |  |  |  |  |
| 8 | 2.10% | 0.60% | 0.50% | -1.50% | -1.65% | 0.12% |
| 17 | 0.30% | 0.00% | 0.00% | -0.30% | -0.23% | -0.03% |
| 14* | 2.40% | 0.40% | 0.00% | -2.00% | 0.00% | 0.00% |
| 18* | 0.30% | 0.07% | 0.00% | -0.20% | 0.00% | 0.00% |
| 12* | 0.80% | 0.00% | 0.00% | -0.80% | 0.00% | 0.00% |
| mean | 97.00% | 100.00% | 98.10% |  |  |  |

## Discussion

KPI was recently evaluated by a group from Genentech as part of a larger effort(16). In a cohort of 72 individuals with ground truth determined by LinkSeq qPCR, Chen et al. report six mismatches in one sample (possibly swapped), and apart from this 95.8% accuracy for *KIR2DS3* and 100% accuracy for the 15 other genes.

This is consistent with the results of our synthetic test, whose accuracy was 100% for all genes. One drawback of the design of the synthetic test is that the haplotypes used in the test were also included in the MSA that was used to generate the per-gene probes. However, the main purposes of the synthetic tests were to test the application of the markers to short reads in a variety of genotypes and recover their original geno- and haplo-types; this is value-added compared with simply demonstrating sequences unique to a gene. The almost-perfect results of the Genentech and synthetic experiments, along with a GoNL results that had a 99.6% family-consistency rate and in line with expected frequencies, provide evidence that KPI’s gene predictions are very accurate.

The evidence also suggests that KPI’s haplotype results are accurate, although often ambiguous: the accuracy in the synthetic test was 100% and the GoNL family-consistency was 99.6, and the predictions allow EM predictions that align with the expected population frequency from the literature.

KIR*IMP’s haplotype frequency estimations differ from expectations in some areas. The evidence from comparisons with frequency reports from Jiang et al. 2012 (Table 5, column 5) suggest KIR*IMP overestimated cA01∼tA01 (haplotype numbers 1 and 2) and underreported haplotypes containing cB01 or cB02 centromeric regions combined with the tA01 telomeric region (cB01∼tA01 and cB02∼tA01, 4, 5, 6, 10) in the GoNL cohort. This discrepancy can also be seen in the predicted genotype frequencies, where KIR*IMP relatively under calls the presence of *KIR2DS2, KIR2DL2*, and *KIR2DS3* by ∼20% and *KIR2DL5* by ∼10% compared with KPI and the historical European frequencies from Allele Frequency Net database (Table 5, column 6); all four of those genes are in cB01, and *KIR2DS2* and *KIR2DL2* are also in cB02. GoNL genotyping was done on the Immunochip, which is the best option according to the KIR*IMP manuscript. With that chip, they report accuracies of 100% for *KIR2DS2*, 98% for *KIR2DL2*, 82% for *KIR2DL5*, 81% for *KIR2DS3*, and 95% for *KIR2DS5* in their Norwegian-German validation cohort. Although the family consistency rate is 100% for KIR*IMP and 96.6% for KPI, their haplotype-pair predictions only agree in 58% of individuals. Without ground truth available, without any reason to expect this cohort to deviate from expectations, and considering KIR*IMP’s self-reported accuracy, the evidence suggests that KPI’s predictions are more accurate than KIR*IMP’s in this cohort and specifically that KIR*IMP under called the presence of genes *KIR2DS2, KIR2DL2, KIR2DS3, KIR2DL5* and haplotypes cB01∼tA01 and cB02∼tA01. There are several potential reasons KIR*IMP’s predictions may be less accurate than KPI’s. The reference haplotypes used for marker discovery for KIR*IMP were defined by copy number genotyping and family relationships; KPI defined its haplotypes using a MSA of full haplotype sequences. KIR*IMP’s input is restricted to a few hundred single nucleotide polymorphisms, whereas KPI can use the entire genomic range of KIR sequences of length 25, which provides the potential for more information per marker and a broader base of markers. KIR*IMP uses a small number of SNPs to call one or more genes, whereas KPI uses dozens-to-thousands of 25mers to call a single gene, One of the steps of KIR*IMP’s workflow is to align and phase all the SNPs to one ‘A’ haplotype, which may be a limitation for genes not on that haplotype; all the gene and haplotypes we found to have lower accuracy rates are not located on the ‘A’ haplotype. KPI has no alignment or assembly steps. It is also important to note that the primary purpose for the comparison with KIR*IMP was not to evaluate the potential success of predicting KIR genes and haplotypes using SNPs vs sequence reads, but rather to compare the two algorithms. Although both algorithms predict the presence/absence of KIR genes and structural haplotypes, their solution domains are very different: microarray SNP panels vs raw genomic DNA reads. Both algorithms report the lowest accuracy rates for *KIR2DS3* and a ∼10% lower frequency rate for *KIR2DS1* in GoNL compared with reference frequencies.

The 85% family consistency rate of the EM-reduced haplotype predictions suggest that KIR haplotype ambiguity cannot be accurately reduced at the individual level via expectation-maximization. However, since the EM-reduced haplotype frequencies are in line with references, it is possible the predictions might aggregate to population-level in a maximum-likelihood manner and therefore perhaps may still be useful for some population genetics purposes.

Traditional lab-based SSO presence/absence genotyping relies on a single short-sequence strategy, an approach that can be applied similarly to synthetic analysis of large amounts of WGS. In this virtual context, primer locations are not needed, and kmer searching is efficient and accurate at populations scales. To develop this synthetic SSO-like (kmer) library, we leveraged the information from a multiple sequence alignment of all full-length haplotypes that are available for this study. We believe this is a more accurate approach than using IPD-KIR reference alleles, because the IPD-KIR reference alleles do not require the haplotype location to be known. In addition, fusion alleles are assigned in IPD-KIR to one of the two parent genes, and therefore large sequences of some alleles are not really from the gene in which they are classified. We used 25 for our ‘k’ (i.e., sequence size) because BLAST searching indicated this to be a conservative minimum length needed to distinguish a small set of test markers to the KIR region. We did not experiment with any k size other than 25, since the choice gives a reasonable number of significant markers and their lack of off-KIR hits as tested the GoNL population WGS confirms the effectiveness as gene/intergenic markers. Having thus obtained the region markers, we then used the most common (‘peak’) hit count from each gene/intergene’s library of sequences to make the PA genotype calls (Supplemental Figure 1). Since KPI decomposes the genetic information into 25mers, it works with any collection of DNA reads, as long as the KIR region is included. It works with fasta, fastq, single, paired, short, and long reads. Since the markers are not unique to exons, it will not work with cDNA or exon only reads.

In this study, we have discovered gene markers and demonstrated their use to predict genes and haplotype-pairs at high accuracy (97%+) with population scale from any kind of sequence data that includes the full KIR region, including WGS. We tested it on synthetic ground-truth sequences and a large cohort of family WGS. We compared our algorithm to the leading SNP-based interpretation algorithm. KPI is free software with a GPL3 license, implemented as a Nextflow workflow backed with an optional Docker environment. It is available at https://github.com/droeatumn/kpi.

## Acknowledgements

This study makes use of data generated by the Genome of the Netherlands Project. A full list of the investigators is available from www.nlgenome.nl. Funding for the project was provided by the Netherlands Organization for Scientific Research under award number 184021007, dated July 9, 2009 and made available as a Rainbow Project of the Biobanking and Biomolecular Research Infrastructure Netherlands (BBMRI-NL). The sequencing was carried out in collaboration with the Beijing Institute for Genomics (BGI). The samples were provided by The LifeLines Cohort Study(17), and generation and management of GWAS genotype data for it, is supported by the Netherlands Organization of Scientific Research (NWO, grant 175.010.2007.006), the Dutch government’s Economic Structure Enhancing Fund (FES), the Ministry of Economic Affairs, the Ministry of Education, Culture and Science, the Ministry for Health, Welfare and Sports, the Northern Netherlands Collaboration of Provinces (SNN), the Province of Groningen, the University Medical Center Groningen, the University of Groningen, the Dutch Kidney Foundation and Dutch Diabetes Research Foundation.

Thanks to Christian Hammer from Genentech for assistance testing and debugging the application. Also, thanks to Cynthia Vierra-Green and Martin Maiers from the Center for International Blood and Marrow Transplant Research (CIMBTR) as well as Julia Udell from Mayo Clinic and the University of Minnesota for KIR consultation and consolation.

## Supporting Information

**Supplemental Figure 1**. Algorithm details. Command to query the markers per genome and details as to how each region was genotyped.

**Supplemental Table 1**. Individual gene and haplotype-pair predictions. Columns A and B contain the family name and relationship. Column C contains KPI’s haplotype-pair predictions, represented by a haplotype list. Each pair is separated with a ‘|’. For example, ‘cA01∼tA01+ cA01∼tB01_2DS5|cA01∼tB01_2DS5+cA01∼tB05’ means the prediction is either haplotype-pairs cA01∼tA01 and cA01∼tB01_2DS5 or haplotype-pairs cA01∼tB01_2DS5 and cA01∼tB05. The haplotype list in column C represents all possible haplotype-pairs fitting the presence-absence genotypes. Column D is a count of the number of haplotype-pair predictions in column C’s haplotype list. Column E is the haplotype list in column C reduced by family relationships, and column F indicates whether or not it is different from the original haplotype list in column C. Column G is the original haplotype list in column C reduced by EM; its results should not be used as they have limited accuracy. Columns H-W are the gene presence-absence predictions.

## Notes

### Competing Interest Statement

The authors have declared no competing interest.

### Summary of Updates

This version includes major changes to the design, implementation, and results.

