## Supplemental Figure 1 for "Accurate and Efficient KIR Gene and Haplotype Inference from Genome Sequencing Reads with Novel K-mer Signatures"

### Algorithm details

#### kmer marker discovery

KPI calls a presence/absence per by aggregating the presence/absence genotype of many small (25mer) probes specific for that gene. This is the command to discover the 25mer  probes for a gene, e.g., *KIR2DL1*

```
kmc -k25 -ci72 -cx72 -fm ./2DL1/2DL1.fasta intersect/2DL1 intersect
kmc_tools complex intersect/./2DL1_cmd.txt
kmc_dump intersect/./2DL1_unique intersect/./2DL1_intersect.txt
```

Once the all probes have been discovered, a KMC database was made for them and their reverse complements

```
kmc -v -k25 -ci1 -fa intersect_union-All_v1_wRc.fasta intersect_union-All_v1_wRc work > intersect_union-All_v1_wRc_err.txt
```

#### Individual kmer genotyping

To genotype a specific individual, a KMC 25mer database was generated from their raw sequences, e.g., 52b. gonl-52b-cmd.txt has six lines, each with the full path to a FASTQ file.

```
kmc -k25 -ci2 -fq @./gonl-52b-cmd.txt ./gonl-52b work
```

Each probes hit count was obtained by interesting the probe database with the individual's genotype database. The output is a collection of marker names and a count for each marker. e.g., 134a

```
kmc_tools -hp simple .//gonl-134a intersect_union-All_v1_wRc intersect gonl-134a -ocleft
kmc_tools dump gonl-134a gonl-134a.txt
```

#### Generating gene presence/absence via probe marker counts

Presence/absence calls for each gene region were made by the peak hit count per region in one individual, with counts of 1 or 2 errors and therefore set to a count of 0. The peak is the most common hit count for all markers in that region. If the most common hit count is zero, the region is called absent; if the hit count is greater than zero, the region is called present.

Figure 1 shows *KIR3DL1*'s 25mer hit distribution for a 'present' genotype in one individual. The most common occurrence (or peak of the chart) is 102 25mers that hit 11 times. Since the peak is greater than 0 hits, *KIR3DL1* is called present.

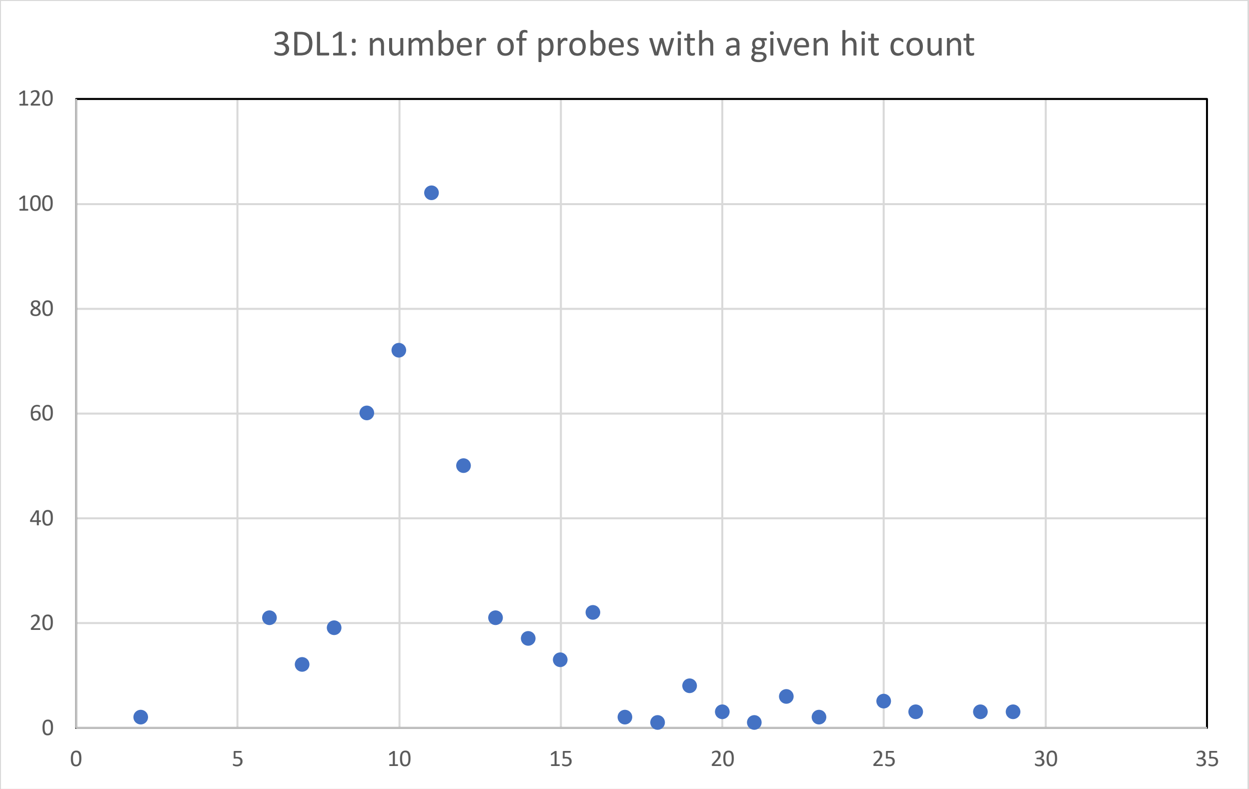


Figure 1. Example hit distribution for a 'present' genotype. The x axis shows the number of hits in the genome. The y axis shows the number of 25mers with that hit count.

Figure 2 shows *KIR3DS1*'s 25mer hit distribution for an 'absent' genotype in one individual. The most common occurrence (or peak of the chart) is 1319 25mers that have 0 hits. Since the peak is 0 hits, *KIR3DS1* is called absent.

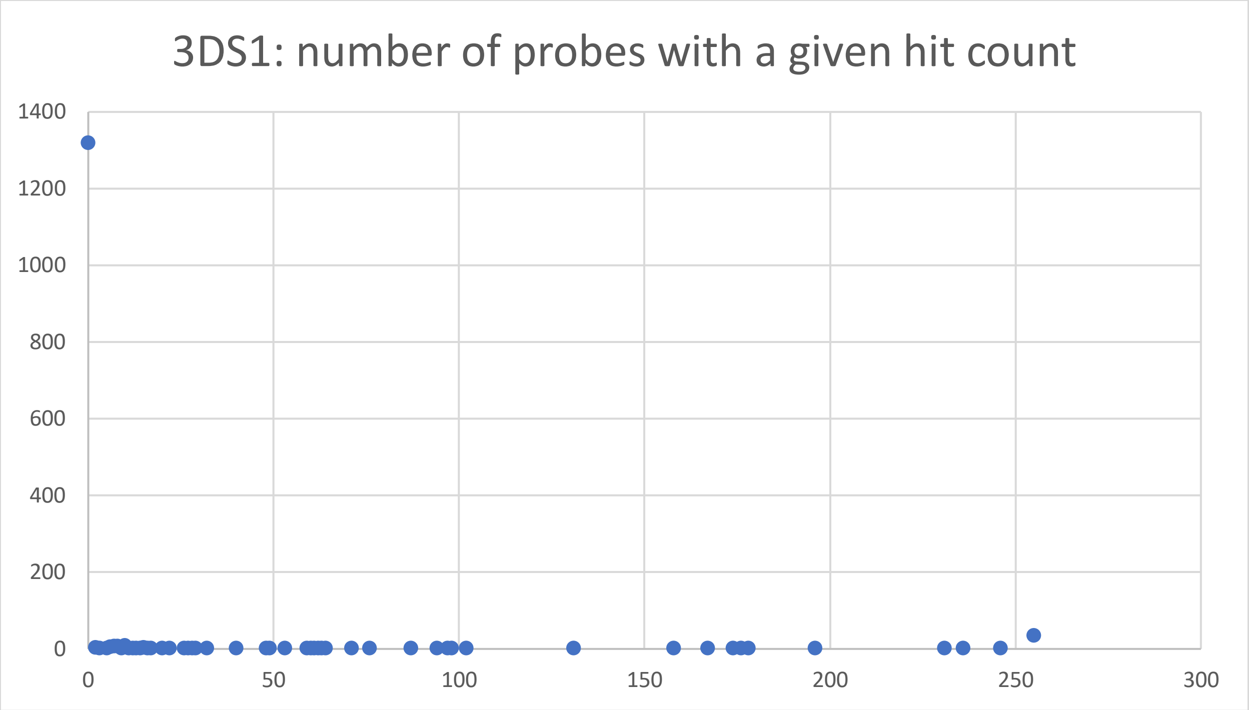

Figure 2. Example hit distribution for a ‘absent’ genotype. The x axis shows the number of hits in the genome. The y axis shows the number of 25mers with that hit count.
